# Employing scan-SCAM to identify residues that compose transmembrane helix 1 (TM1) of EnvZ

**DOI:** 10.1101/2020.08.25.266015

**Authors:** Thanwarat Chavalwan, Fadhael Alrahman H. Hasan, Phillipa Cheesman, Rahmi Yusuf, Roger R. Draheim

## Abstract

The *Escherichia coli* sensor kinase EnvZ modulates porin expression in response to various stimuli including intracellular osmolarity, intracellular pH and periplasmic interaction with MzrA. The expression of two major outer membrane porins, OmpF and OmpC, are regulated by EnvZ, and act as passive diffusion-limited pores allowing compounds, including certain classes of antibiotics such as β-lactams and fluoroquinolones, to enter the bacterial cell. Even though allosteric processing occurs within both the periplasmic and cytoplasmic domains of EnvZ, how the transmembrane domain bi-directionally transmits these signals remains not fully understood. Here, we employ a library of single-Cys-containing EnvZ proteins to perform scan-SCAM in order to map the precise residue composition of TM1. Our results demonstrate that residue positions 19 through 30 reside within the membrane core and compose a tightly packed portion of TM1. We also show that positions 15 through 18 and position 31 are interfacial and slightly splayed apart compared to those tightly packed within the hydrophobic core. Finally, we reveal that residue positions 33 and 34 reside in the periplasm and participate in robust protein-protein interactions, while the periplasmic positions 35 through 41 exhibit helical periodicity. We conclude by synthesizing these new insights with recent high-resolution structural information into a model of membrane-spanning allosteric coupling between the periplasmic and cytoplasmic domains of EnvZ.

## 1 Introduction

Recent modelling demonstrates that when antimicrobial resistance (AMR) remains left unchecked, 300 million people are predicted to expire prematurely by 2050 and 100 trillion USD worth of economic output will be lost [1]. Within this context of AMR, Gram-negative organisms are particularly troublesome as most antibacterial chemotherapeutics need to pass through a lipopolysaccharide (LPS)-coated outer membrane (OM) and those that gain entry are often pumped back out by efflux pumps [2, 3]. The OM serves as the first line of defence for Gram-negative bacteria as it is impermeable to large, charged molecules and influx is controlled by porins [3–5]. These porins are the major pathway for prominent groups of antibiotics, such as beta-lactams and fluoroquinolones into the cell interior [6]. Furthermore, given their role in antibiotic uptake, it should not be surprising that porin expression is often changed in patients undergoing antibiotic treatment [7–13].

The EnvZ/OmpR two-component system (TCS) is present within many Gram-negative bacteria and was initially shown to respond to changes in external osmolarity by modulating porin expression within the outer membrane [3, 14]. Two of the major porins expressed within the outer membrane of *Escherichia coli* are OmpF and OmpC, a large- and small-diameter porin, respectively and are governed by EnvZ signal output. When the EnvZ/OmpR TCS is activated, more OmpC than OmpF is present in the outer membrane and this significantly reduces the permeability of the cell membrane [3]. Over the last few years, responses to three stimuli have been studied in greater molecular and biophysical detail than previously possible: changes in intracellular osmolarity, changes in intracellular pH and periplasmic interactions with MzrA (Figure 1). Recent biophysical analysis has resulted in a “stretch-relaxation” model proposing that increased intracellular osmolarity promotes a more folded conformation of a cytoplasmic four-helix bundle of EnvZ due to increased stabilisation of intra-helical hydrogen bonding [15, 16]. This conformation allows enhanced rates of autophosphorylation of the conserved His-243 residue and subsequent phosphotransfer to OmpR. In another study, intracellular pH was shown to dramatically affect the phosphatase activity of EnvZ [17]. Finally, MzrA (<u>m</u>odulator of Env<u>Z</u> and Omp<u>R</u> protein <u>A</u>) was initially identified and characterised as a small inner membrane “connector” protein that interacts with the periplasmic domain of EnvZ *in vivo*. MzrA-EnvZ interactions have been shown to result in increased EnvZ signal output in an osmosensing-independent manner [18, 19].

**Figure 1.**
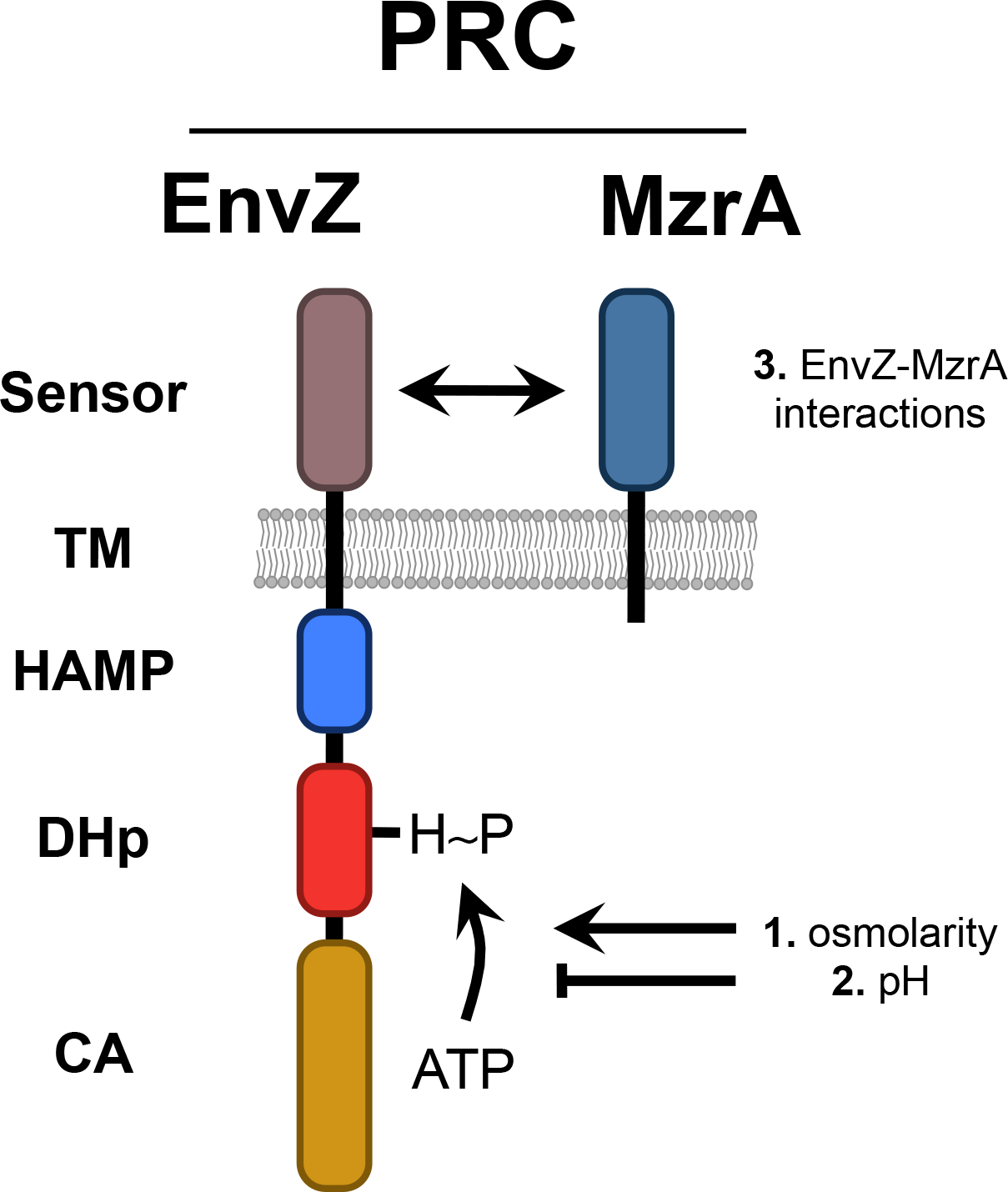
The porin regulatory complex (PRC) is composed of EnvZ and MzrA. Recently, the effects of three stimuli on modulation of PRC signal output have been examined in great biochemical/biophysical detail: the effect of osmolarity on the cytoplasmic domain of EnvZ [15, 16], the effect of pH on the cytoplasmic domain [17] and interactions between EnvZ and MzrA in the periplasm [18, 19].

Based on these studies, we desired to generate a more complete understanding of the organisation of the TM domain of EnvZ and its role during stimulus perception and processing. We began by using a previously created library of EnvZ receptors that each contains a single Cys substitution between residues positions 11 and 41 to perform scan-SCAM. Our results demonstrate that residue positions 19 through 30 are tightly packed and embedded within the lipid core of the inner membrane, while positions 15 through 18 and position 31 remain interfacial and slightly splayed apart. On the periplasmic end of TM1, positions 33 and 34 are likely engaged in robust protein-protein interactions whereas residue positions 35 and onward reside in the periplasm and adopt a helical periodicity. We conclude by discussing these results within the context of the other recently obtained structural information and propose a model for transmembrane communication by EnvZ.

## 2. Materials and Methods

### 2.1 Bacterial strains and plasmids

*E. coli* strain MC1061 [20] was used for all DNA manipulations, while strain EPB30 [21], which is a *envZ::kan* derivative of MDG147 [MG1655 ϕ(*ompF^+^*-*yfp^+^*) ϕ(*ompC^+^*-*cfp^+^*)] [22] was used for all Cys-reactivity experimentation. Plasmid pRD400 [23] retains the IPTG-based induction of EnvZ originally present in plasmid pEnvZ [24] while also encoding a seven-residue linker (GGSSAAG) [25] and a C-terminal V5 epitope tag (GKPIPNPLLGLDST) [26]. Standard molecular biological and microbiological techniques were employed for plasmid DNA extraction (QIAprep Spin Miniprep Kit, QIAgen, Venlo, Netherlands), confirmation of Cys residue position via plasmid-based sequencing (Eurofins Genomics, Ebersberg, Germany) and transformation of sequence-confirmed plasmids [27].

### 2.2 Selection of residues for assessment

The selection of residues previously assessed to contain TM1 was described in [28]. Briefly, the primary sequence of EnvZ from *Escherichia coli* K-12 MG1655 (NP_417863.1) was subjected to a full protein scan with DGpred, which calculates the Δ*G_app_* for transmembrane insertion throughout the entire length of the submitted sequence [29]. NP_417863.1 (EnvZ) was also assessed for putative transmembrane helices by a hidden Markov model with TMHMM v2.0 [30]. Both software packages highlighted a motif commonly found within TM helices consisting of positively charged residues with adjacent aromatic residues that bracket a core of aliphatic residues as a potential candidate for TM1 [31]. Based on these results, we previously created a single-Cys-containing library of plasmid-based EnvZ variants that spanned each individual position from 11 to 41, with the exception of an L32C variant that could not be constructed [28].

### 2.3 Cys-reactivity experimentation

We modified a previously employed scan-SCAM protocol [32] to function with our plasmid-based expression system of single-Cys-containing EnvZ variants. Briefly, EPB30/pRD400 cells were grown as described previously [23] with slight modifications. Fresh colonies were used to inoculate 2-ml overnight cultures of lysogeny broth [33]. Cells were grown overnight at 37 °C and diluted at least 1:1000 into 3 ml of fresh minimal medium A [33] supplemented with 0.2% glucose. Ampicillin, sucrose and IPTG were present where appropriate. Upon reaching an OD_600nm_ ~ 0.6, 1.33 ml of cells were harvested by centrifugation at 17000 x *g* for 15 min at room temperature. After centrifugation, cells were washed once with 50 mM HEPES containing 600 mM NaCl, 4 mM MgSO_4_ and 1 mM EDTA at pH 6.8 and again with 50 mM HEPES at pH 6.8 at which time they were pelleted by centrifugation anew and subjected to one of three different treatment regimens. For the SP treatment, cells were resuspended in 50 mM HEPES containing 4% SDS at pH 6.8 and subjected to one freeze/thaw cycle to aid in solubilisation of the cell membrane. PEG-Mal was then added to a final concentration of 5 mM and samples were incubated in the dark at room temperature for 1.5 hours with intermittent vortexing for 10 seconds every 15 min. SDS-PAGE buffer containing 250 mM DTT was added to stop the reaction. For the SNP treatment, cells were resuspended in 50 mM HEPES containing 4% SDS at pH 6.8 and subjected to one freeze/thaw cycle to aid in solubilisation of the cell membrane. *N*-Ethylmaleimide (NEM) was then added to a final concentration of 2 mM and samples were incubated in the dark at room temperature for 1.5 hours with intermittent vortexing for 10 seconds every 15 min. PEG-Mal was then added to a final concentration of 5 mM and samples were incubated in the dark at room temperature for 1.5 hours with intermittent vortexing for 10 seconds every 15 min. SDS-PAGE buffer containing 250 mM DTT was added to stop the reaction. Finally, for the NSP treatment, cells were resuspended in 50 mM HEPES buffer at pH 6.8 and NEM was added to a 2mM final concentration and samples were incubated in the dark at room temperature for 1.5 hours with intermittent vortexing (every 15 min for 10 seconds.) Samples were then washed twice with 50 mM HEPES buffer at pH 6.8 and resuspended in 50 mM HEPES containing 4% SDS at pH 6.8 and subjected to one freeze/thaw cycle to aid in solubilisation of the cell membrane. PEG-Mal was then added to a final concentration of 5 mM and samples were incubated in the dark at room temperature for 1.5 hours with intermittent vortexing for 10 seconds every 15 min. SDS-PAGE buffer containing 250 mM DTT was added to stop the reaction.

### 2.4 Analysis of Cys-reactivity samples via SDS-PAGE and immunoblotting

Samples were subsequently analysed on 7.5% SDS/acrylamide gels. Standard buffers and conditions were used for electrophoresis, immunoblotting and detection with enhanced chemiluminescence [34]. Anti-V5 (Invitrogen) was used as the primary antibody, while peroxidase-conjugated anti-mouse IgG (Sigma) was employed as the secondary antibody. Digitized images were acquired with a ChemiDoc MP workstation (Bio-Rad). When no slower-migrating PEGylated band was observed with a particular combination of Cys residue position and experimental condition, a second independent experiment was performed. When a slower-migrating band was observed, a minimum of three independent experiments were performed. ImageJ v1.49 [35] and QtiPlot v0.9.8.10 (IONDEV SRL, Bucharest, Romania) were used as previously described to quantify band intensities [28].

## 3 Results

### 3.1 Description of the various Cys-reactivity treatments

To retain the functional integrity of the EnvZ/OmpR osmosensing circuit, we determined the baseline membrane-relative position of EnvZ TM1 *in vivo*. A scan-SCAM method [27, 32], which employs a single-Cys-containing library of EnvZ receptors allowed us to precisely map the residues that compose TM1 and how they might reposition when cells were grown under a high-osmolarity (15% sucrose) regime. To employ scan-SCAM, we used a previously made Cys-less variant of EnvZ (C277A) and a previously made library of single-Cys-containing variants over the amino acid position range of 11 to 41 [28]. Scan-SCAM employs various treatments that involving combinations of blocking with *N*-ethylmaleimide (NEM; N), solubilisation of the bacterial cell with sodium dodecyl sulfate (SDS; S) and modification with the thiol-reagent methoxypolyethylene glycol maleimide (PEG-mal; P) that possesses a long polyethylene glycol chain of approximately 5 kDa. Modification with PEG-mal facilitates observation of slower-migrating band via SDS-PAGE and immunoblotting against EnvZ possessing a C-terminal V5-epitope tag.

### 3.2 Mapping residues composing TM1 of *E. coli* EnvZ *in vivo*

We began by performing the SP treatment with EPB30/pRD400 cells expressing one of the single-Cys-containing EnvZ variants from the TM1 library. We grew cells under both the low-osmolarity (0% sucrose) and high-osmolarity (15% sucrose) regimes to highlight any differences in TM1 positioning. The SP treatment involves solubilisation of bacterial cells in buffer containing 4% SDS and subsequent treatment with PEG-mal. Following treatment, lysates were subjected to SDS-PAGE and immunoblotting to identify slower-migrating PEG-lyated bands (Figure 2). Similar patterns of PEG-lyation were observed whether cells were grown under the low- or high-osmolarity regimes. However, slight differences were seen within residues exposed to the cytoplasm or periplasm.

**Figure 2.**
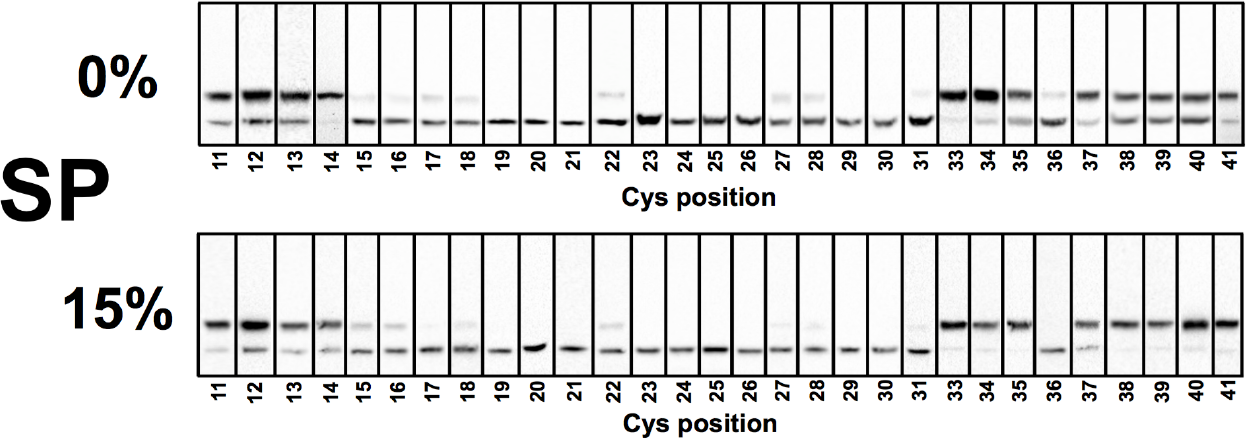
SP treatment of EPB30/pRD400 cells expressing one of the single-Cys-containing EnvZ receptors. Cells expressing one of the single-Cys-containing EnvZ variants were grown under the low-(0% sucrose) or high-osmolarity (15% sucrose) regime. Under both regimes, residue positions 11 to 14 and 33 to 41, with the exception position of 36, were subject to high levels of PEG-ylation appearing as a slower-migrating band. Residue positions 15 to 18, 22, 27, 28, 31 and 36 were subject to low levels of PEG-ylation. Other residue positions exhibited no PEG-ylation.

In both cases, a greater extent of PEG-ylation was observed between residue positions 11 and 14 and at most residue positions between 33 and 41. A lesser extent of PEG-ylation was observed between residues positions 15 and 18 and at residues positions 22, 27-28, 31 and 36. At least three samples were collected for each residue position, analysed and subjected to quantification as shown in Figure 3. We also subjected EPB30/pRD400 cells grown under the low- or high-osmolarity regimes to the SNP treatment to confirm that the sulfhydryl groups present on the Cys residues were blocked by NEM. Regardless of growth regime, we observed that all free PEG-ylatable sulfhydryl groups under the SP condition were blocked by NEM (Figure 4). In summary, our data suggests that residues 19 through 30 composed a tightly packed hydrophobic core of TM1 from EnvZ *in vivo*. Positions 15 through 18 and 31 likely resided within the polar/hydrophobic interfacial regions of the membrane and remain slightly splayed apart.

**Figure 3.**
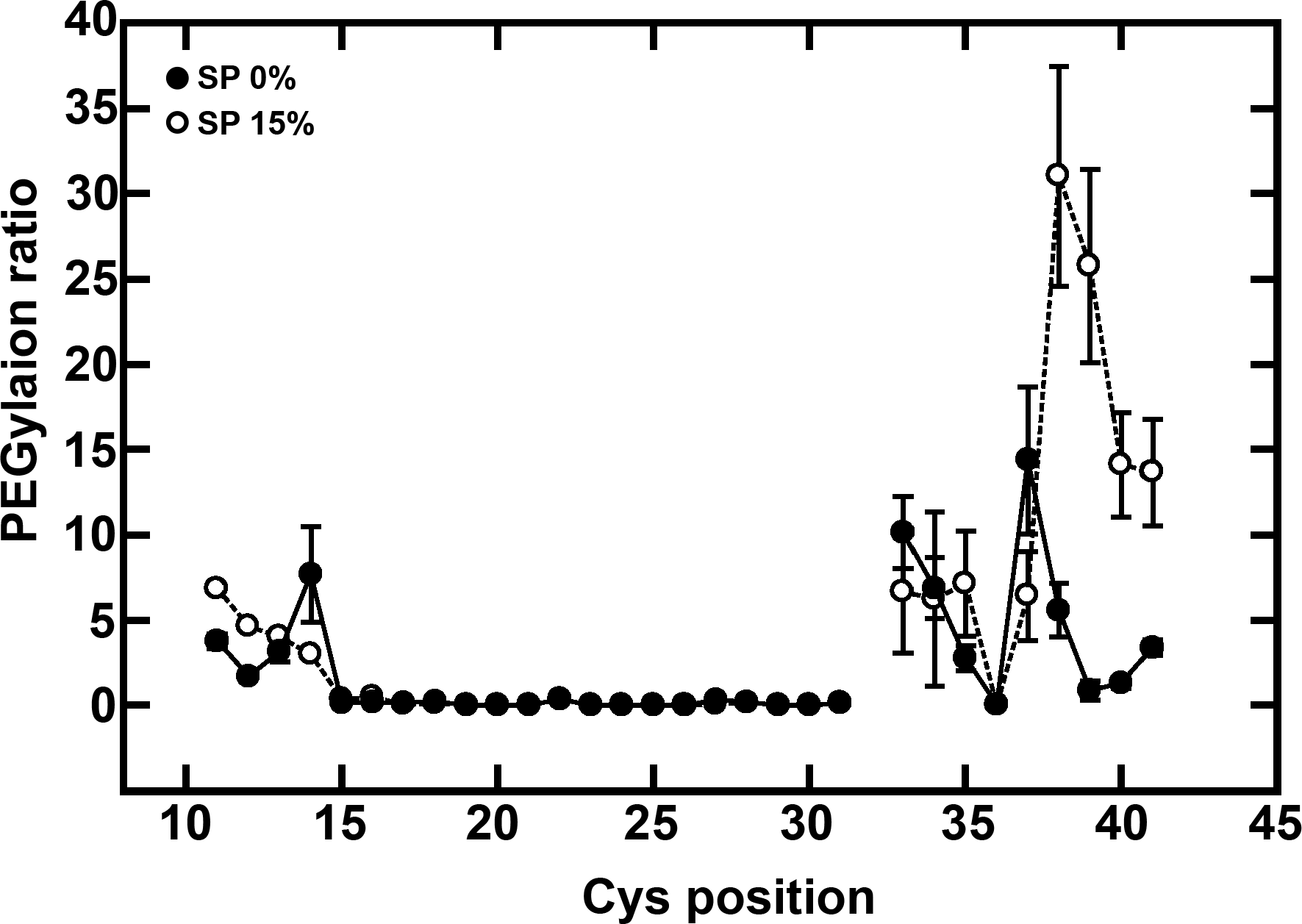
Quantification of PEG-ylation. Residue positions 11 to 14 exhibit high levels of PEG-ylation and quantified by the ratio of PEG-ylated to un-PEG-ylated moieties present in at least 3 samples. Positions 15 through 31 are PEG-ylated to a minor extent, if at all, while positions 33 through 41 exhibit a helical pattern of PEG-ylation. Where large amounts of PEG-ylation occur, EPB30/pRD400 cells exhibit greater PEG-ylation when grown under the high-osmolarity (15% sucrose) regime.

**Figure 4.**
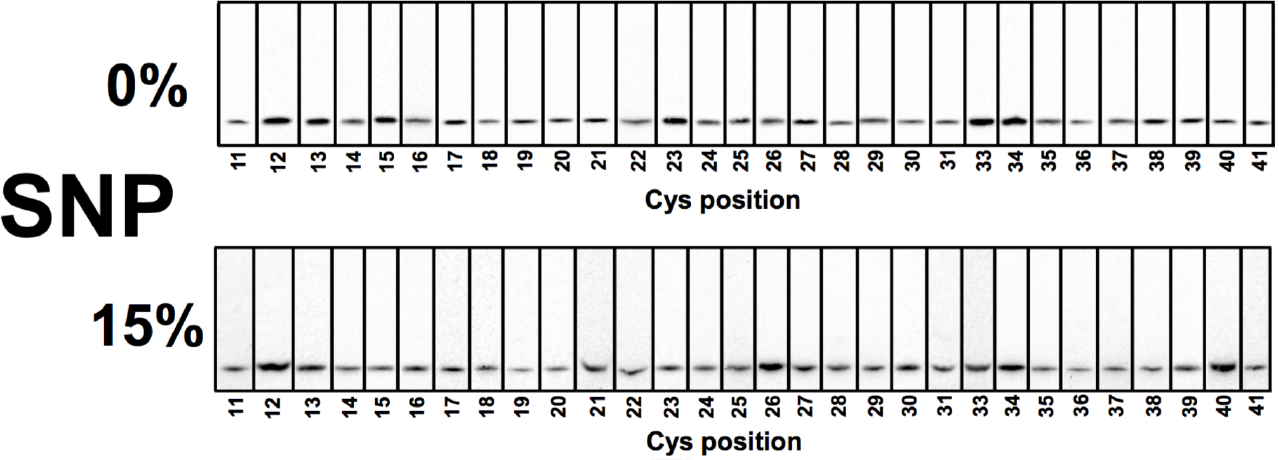
NSP treatment of EPB30/pRD400 cells expressing one of the single-Cys-containing EnvZ receptors. Cells were grown under the low-(0% sucrose) or high-osmolarity (15% sucrose) regime expressing one of the single-Cys-containing EnvZ variants. Under both regimes, all PEG-ylatable residue positions were blocked by NEM.

### 3.3 Mapping the solvent-exposed residues of EnvZ TM1

After mapping TM1 *in vivo*, we employed the NSP treatment, which facilitates blocking of solvent-exposed Cys residues with NEM prior to PEG-ylation. Comparison between results obtained with the SP and NSP treatment allowed us to determine whether any residues within TM1 were solvent-exposed (Figure 5). All residues positions between 11 and 21 were blocked by NEM, as were residue positions distal to position 34. Residue positions 22, 27-28, 31 (at 15% sucrose) and 33-34 remain susceptible to PEG-ylation and thus demonstrate that blocking by NEM was not complete. In addition, the pattern of PEG-ylation remains similar regardless of the growth regime employed to grow EPB30/pRD400 cells (Figure 6).

**Figure 5.**
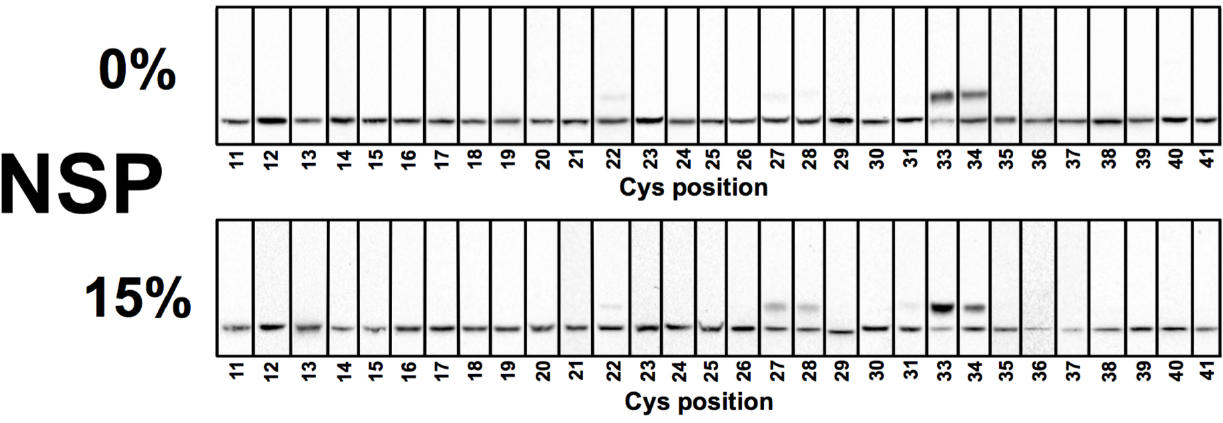
SNP treatment of EPB30/pRD400 cells expressing one of the single-Cys-containing EnvZ receptors. Cells were grown under the low-(0% sucrose) or high-osmolarity (15% sucrose) regime. Under both regimes, PEG-ylation that was previously observed at residue positions 11-21 was blocked by NEM. However, PEG-ylation at residue positions 22, 27-28, 31 (with 15% sucrose) and 33-34 remained. All residue positions from 35 onward were completely blocked by NEM.

**Figure 6.**
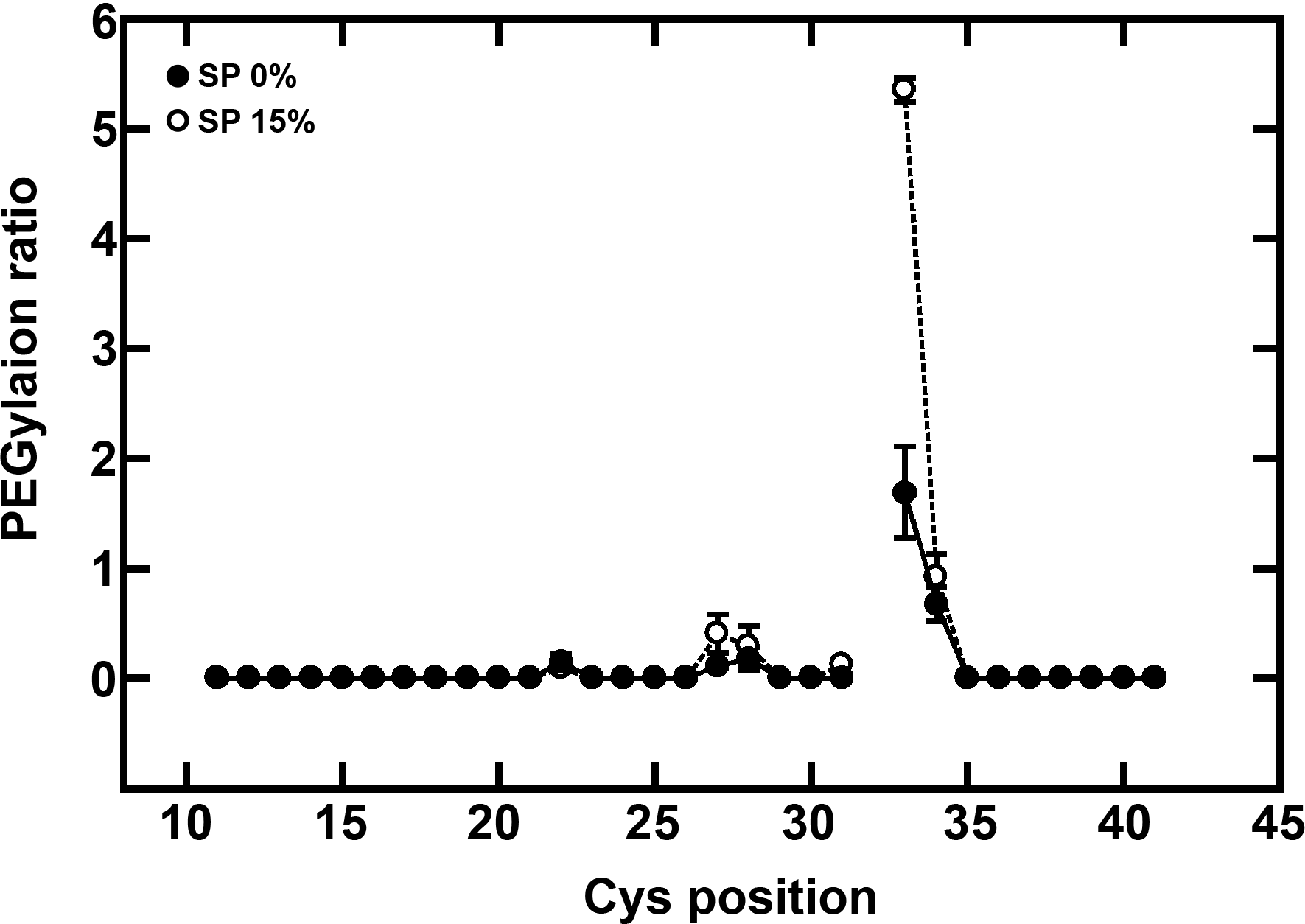
Quantification of PEG-ylation after blocking with NEM. Residue positions 22, 27 and 28 are subject to minimal levels of PEG-ylation that are similar to the SP treatment. Residue positions 33 and 34 remain PEG-ylatable as well, suggesting that they are in a tight protein-protein contact that prevents blocking by NEM.

## 4 Discussion

### 4.1 Delineation of TM1 boundaries

Here, we sought to delineate the boundaries of TM1 from *E. coli* EnvZ *in vivo*. We submitted the *E. coli* K-12 MG1655 EnvZ primary sequence (NP_417863.1) to two software packages, DGpred [29] and TMHMM v2.0 [30], which proposed that residue positions 15-37 and 12-34 compose TM1, respectively. These predictions suggested that we could use our pre-existing single-Cys-containing EnvZ library to determine the boundaries. Previous work demonstrated the membrane topology of EnvZ and the presence of TM1 and predicted that residues 16 through 41 or 16 through 46 passed through the membrane [36, 37].

Our analysis with the SP treatment demonstrated that residue positions 19 through 30 of *E. coli* EnvZ were tightly packed and located within the bilayer core as these residues were not subject to high levels of PEG-ylation (Figures 2, 3 and 7.) Unfortunately, we were not able to substitute the leucine residue present at EnvZ position 32 for a cysteine residue, hinting at its critical role in EnvZ function or stability, and thus, the precise location of this residue position remains unclear. This range of residues, 19 through 30, agrees with our previous *in vivo* sulfhydryl-reactivity experimentation demonstrating that positions 19, 23, 26 and 30 are in close proximity to each other and form the TM1-TM1’ dimeric interface within EnvZ *in vivo* (Figure 7) [28].

**Figure 7.**
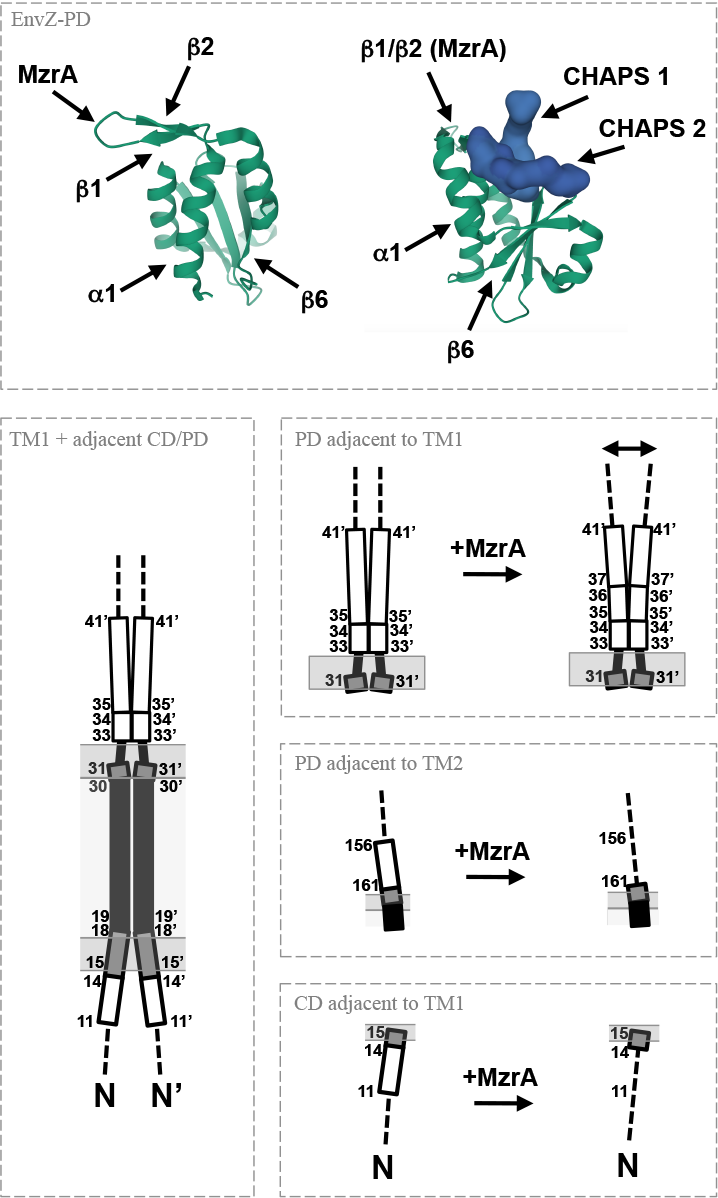
Proposed mechanism of α-helicity modulation during TM communication by EnvZ. A recent high-resolution structure of the periplasmic domain from *E. coli* EnvZ (EnvZ-PD) demonstrates both a CHAPS-binding cleft and a β-hairpin required for MzrA interaction [41]. Here we demonstrate that modulation of helicity in the cytoplasm and periplasmic residues adjacent to TM1 occurs in a growth regime-dependent manner. This newly gleaned detail can be merged with previously identified modulation of helicity in the periplasmic residues adjacent to TM2 to formulate testable models for transmembrane communication by EnvZ.

### 4.2 Osmolarity-dependent structures at the cytoplasmic and periplasmic ends of TM1

When cells expressing the single-Cys-containing library were subjected to the NSP treatment, residue positions 11 through 18, which were previously accessible to PEG-ylation under the SP treatment regime, were blocked by NEM (Figures 2–3 and 5–6). This demonstrated that while residue positions 15 through 18 were located within the membrane and rendering them difficult to access by PEGmal possessing an approximately 5 kDa moiety, the small NEM molecule was able to block the interfacial residues at these positions, suggesting that the cytoplasmic end of the TM bundle is as not tightly packed as it is within the hydrophobic core (Figure 7). At low omolarity, residue position 31 exhibits similar character at the periplasmic end of TM1 (Figures 2–3 and 5–6). Additionally, evidence of helical formation in residues 11 through 14 was present when cells are grown under the low-osmolarity regime and lost when the cells were grown under the high-osmolarity regime (Figures 2, 3 and 7). Through the membrane core, residue positions 22, 27 and 28 remain unblocked by NEM and subject to PEG-ylation after SDS treatment (Figure 4).

At periplasmic end of TM1, blocking by NEM occurred on residues distal to position 35. In contrast, residue positions 33 and 34 were highly PEG-ylatable after SDS treatment but not blockable with NEM prior to SDS treatment. One possible explanation is that residue positions 33 and 34 remain PEG-ylatable after disruption of protein-protein interactions with SDS during SP treatment because they do not reside within the membrane core. However, due to potential robust protein-protein interactions, NEM cannot block these residues prior to disruption with SDS, leaving them subject to PEG-ylation during the NSP treatment. In a similar manner to the cytoplasmic-based positions, differences when cells were grown at the low- and high-omsolarity regimes were also observed distal to the periplasmic end of TM1. Both sets exhibit roughly helical character, but residues positions were subject to greater PEG-ylation with cells grown at higher osmolarity (Figure 7).

### 4.3 Allosteric coupling within the EnvZ-MzrA porin regulatory complex (PRC)

When the results presented here are merged with previous structural information from EnvZ and MzrA, a more detailed understanding of how the composite porin regulatory complex (PRC) functions becomes apparent (Figure 7). Initial topological analysis determined that N-terminus of EnvZ resides within the cytoplasm with a residue composition consistent with the positive-inside rule, while TM1 functions as a signal sequence [37–39]. Here, we detailed that residues 11 through 14 of EnvZ were located within the cytoplasm with a helical structure when the cells were grown at low osmolarity and more extended and exposed to solvent when grown under the high-osmolarity regime (Figures 2 and 3). Previous studies with MzrA have shown that it adopts a topology consisting of an N-terminal cytoplasmic tail, a single TM helix and a periplasmic C-terminal domain [18]. The recent high-resolution structure of the combined four-helix-bundle/TM/HAMP domain of *E. coli* NarQ illustrates significant interaction between the N-terminal cytoplasmic tail and TM1 with the cytoplasmic HAMP domain [40]. Although not essential for MzrA-based modulation of EnvZ signal output [18, 19], it appeared that when cells were grown under the high-osmolarity regime, which results in the presence of MzrA, the EnvZ residues cytoplasmic to TM1 lost their α-helicity and became more solvent-exposed (Figures 2 and 3). These results suggest that a structural rearrangement occurs within the lengthy cytoplasmic tail of EnvZ, which might also modulate signal output in a similar manner to *E. coli* NarQ (Figure 7). *In vivo* experimentation is currently underway to better understand these and other intra-protein and inter-protein couplings within the porin regulatory complex (PRC).

Residue positions 15 through 18 and position 31 of EnvZ reside within the membrane-water interfacial region, as evidenced by a reduced extent of PEG-ylation (Figures 2 and 3), and remained splayed apart as shown via blocking with NEM during the NSP treatment regime (Figures 5 and 6). Positions 19 through 30 were embedded within the hydrophobic core of the membrane, based on very low levels of PEG-ylation under the SP regime and lack of NEM-based blocking during the NSP treatment, which is in agreement with previous results demonstrating that residues 19, 23, 26 and 30 interact with each other and form the TM1-TM1’ interface (Figures 2–3 and 5–6). Unfortunately, position 32 could not be mapped as it was Cys-intolerant. However, the adjacent positions 33 and 34 were found to be involved in robust protein-protein interactions, while the remainder of the periplasmic residues examined, those through position 41, exhibit a helical periodicity based on the extent of PEGylation observed under the SP treatment (Figures 2–3). However, once more, the solvent exposure was greater at high osmolarity, suggesting that a conformational change was occurring here as well (Figure 7).

A high-resolution structure of the periplasmic domain of *E. coli* EnvZ (EnvZ-PD) encompassing residues 36 through 158 was recently determined that consists of a mixed α/β structure with a central β-sheet flanked by α-helices on both sides [41] (Figure 7). Within this structure, the first α-helix (α1) forms after Pro-41, which is the residue most distal to the membrane that we analysed. Therefore, we propose that TM1-α1 could possibly form an extended helix structure possessing three residues, Leu-43, Leu-50 and Leu-57, which have been shown to participate in a functionally important leucine zipper [36]. This extended helix leads into a hairpin-like β-strand composed of β1 and β2, terminating with a poly-Pro motif from residues 71 through 75 (Val-Val-Pro-Pro-Ala) that inhibits homodimerisation of the purified EnvZ-PD [41]. This VVPPA poly-Pro motif has been recently shown to be functionally important for interaction with MzrA and resultant modulation of EnvZ signal output [18, 19, 42]. A related structure of the periplasmic domain of *Salmonella enteritica* serovar Typhimurium PhoQ (PhoQ-PD) possesses a similar hairpin-like β-strand composed of β1 and β2 [43]. Both EnvZ and PhoQ interact with a small biotopic protein, MzrA and MgrB respectively, that consists of a cytoplasmic N-terminus, a single TM helix and a small periplasmic domain suggesting correlations could be drawn between the presence of this hairpin-like β-strand and the existence of membrane-embedded interaction partners [18, 19, 44] (Figure 7).

A central β-sheet composed of antiparallel strands β3 through β6 and several α-helices, namely α1 α2 and α4 form a binding cleft where two CHAPS molecules are bound (Figure 7). Based on competition experimentation and 2D ^1^H-^15^N TROSY spectral analysis, it is proposed that sodium cholate, a biologically relevant bile salt with a similar structure to CHAPS, binds in this same cleft. It was also demonstrated that CHAPS and sodium cholate modulate EnvZ output suggesting that bile salts modulate porin balance via an EnvZ-based mechanism within the human gut [41]. When the CHAPS-bound periplasmic EnvZ domain (EnvZ-PD) is compared to *apo* structure from similar periplasmic domains, a significant change in orientation of the β6 strand is observed [41]. Within the EnvZ-PD structure, terminal to this final β-strand (β6) is a proline residue (Pro-148) and an unstructured tail.

Previous sulfhydryl-reactivity experimentation has revealed that the TM2-TM2’ interface is composed of residues 164, 167, 171, 175 and 179 and that minimal change in disulphide-formation is observed when cells grown under the low- or high-osmolarity regime are compared [45]. However, a growth-regime-dependent difference in sulfhydryl-reactivity at residue positions 156, 157, 160 and 161 has been observed. At low osmolarity, and thus in the absence of MzrA, a helical pattern of cross-linking is observed between positions 156 and 161 that is abolished when cells are grown under the high-osmolarity regime, or in the presence of MzrA [45], which strongly suggests that modulation of helicity might be responsible for transmembrane communication (Figure 7). In the *apo* structure of *E. coli* NarQ, a continuous helical transition occurs from to TM1 to α1, whereas in the *holo* structures, a break is the continuous helix is present [40]. This could be similar to what is observed at the periplasmic ends of TM1 and TM2 in EnvZ. On the opposite end of TM2, these changes could be transmitted downstream via a control cable serving as a physical coupler between the cytoplasmic end of TM2 and the HAMP domain [46–48], where the AS1 sequences from various bacterial receptors have been shown to possess differing propensities for α-helicity [49]. This control cable connects to the HAMP domain [48, 50, 51] and allosterically couples changes in the TM domain to the cytoplasmic domains responsible for signal output of the PRC.

### 4.4 Conclusion

When the information presented here is synthesised with the other previously existing datasets from EnvZ and MzrA, a clearer picture of signal transmission based upon modulation of α-helicity of the membrane-adjacent domains within the porin regulatory complex (PRC) becomes possible. We are currently undertaking a rigorous analysis of EnvZ-MzrA and MzrA-MzrA interactions *in vivo* and building high-resolution models for *in silico* screening in order to identify potential modulators of PRC signal output. It has recently come to our attention that an analogous approach is currently being undertaken with respect to the membrane-embedded PhoQ/MgrB complex of *E. coli* lending credibility to this long-term strategy [44].

## Acknowledgements

R. Y. was generously supported by the Indonesia Endowment Fund for Education, Ministry of Finance (S-4833/LPDP.3/2015), while F. H. H. was sponsored by the Kuwaiti Civil Service Commission Scholarship Program - Ministry of Health. This work was also supported with start-up funding from the Faculty of Science and from the Institute of Biomedical and Biological Science (IBBS) to R. R. D. at the University of Portsmouth. Monabel May Birao, Peltine Ndifon, and Joy Yau (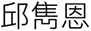) rigorously proofread and provided useful commentary on various iterations of the manuscript that resulted in its improvement.

## Abbreviations

AMR: antimicrobial resistance
AS1: amphipathic sequence 1
CD: cytoplasmic domain
CHAPS: 3-((3-cholamidopropyl) dimethylammonio)-1-propanesulfonate
Cys: cystyl residue
DNA: deoxyribonucleic acid
DTT: dithiothreitol
HAMP: domain found in <u>h</u>istidine kinases, <u>a</u>denylate cyclases, <u>m</u>ethyl-accepting chemotaxis proteins and <u>p</u>hosphatases
HEPES: (4-(2-hydroxyethyl)-1-piperazineethanesulfonic acid), His, histidyl residue
IgG: immunoglobulin G
IPTG: isopropyl-β-D-1-thiogalactopyranoside
LPS: lipopolysaccharide
kDa: kilodalton
NEM: *N*-ethylmaleimide
OD_600nm_: optical density at 600nm
OM: outer membrane
PD: periplasmic domain
PEG: polyethylene glycol
PEG-mal: methoxypolyethylene glycol maleimide
PRC: porin regulatory complex
SCAM: substituted cysteine accessibility method
SDS: sodium dodecyl sulphate
SDS-PAGE: sodium dodecyl sulphate – polyacrylamide gel electrophoresis
TM: transmembrane
TM1: transmembrane helix 1
TM2: transmembrane helix 2
TCS: two-component system
TROSY: transverse relaxation-optimized spectroscopy
USD: United States dollars

## Notes

### Competing Interest Statement

The authors have declared no competing interest.

## References

[1] J. O'Neill, Review on Antimicrobial Resistance. Antimicrobial resistance: tackling a crisis for the health and weath of nations, in, Review on Antimicrobial Resistance, 2014, pp. 20.

[2] H. Nikaido, Prevention of drug access to bacterial targets: permeability barriers and active efflux, Science, 264 (1994) 382–388.

[3] H. Nikaido, Molecular basis of bacterial outer membrane permeability revisited, Microbiology and molecular biology reviews: MMBR, 67 (2003) 593–656.

[4] A.H. Delcour, Solute uptake through general porins, Frontiers in bioscience: a journal and virtual library, 8 (2003) d1055–1071.

[5] G.E. Schulz, The structure of bacterial outer membrane proteins, Biochim Biophys Acta, 1565 (2002) 308–317.

[6] A. Bryskier, Antimicrobial Agents: Antibacterials and Antifungals, in, ASM Press, Washington, DC, USA, 2005.

[7] E. Elliott, A.J. Brink, J. van Greune, Z. Els, N. Woodford, J. Turton, M. Warner, D.M. Livermore, In vivo development of ertapenem resistance in a patient with pneumonia caused by Klebsiella pneumoniae with an extended-spectrum beta-lactamase, Clinical infectious diseases: an official publication of the Infectious Diseases Society of America, 42 (2006) e95–98.

[8] S. Hernandez-Alles, M. Conejo, A. Pascual, J.M. Tomas, V.J. Benedi, L. Martinez-Martinez, Relationship between outer membrane alterations and susceptibility to antimicrobial agents in isogenic strains of Klebsiella pneumoniae, The Journal of antimicrobial chemotherapy, 46 (2000) 273–277.

[9] G.A. Jacoby, D.M. Mills, N. Chow, Role of beta-lactamases and porins in resistance to ertapenem and other beta-lactams in Klebsiella pneumoniae, Antimicrob. Agents Chemother., 48 (2004) 3203–3206.

[10] F.M. Kaczmarek, F. Dib-Hajj, W. Shang, T.D. Gootz, High-level carbapenem resistance in a Klebsiella pneumoniae clinical isolate is due to the combination of bla(ACT-1) beta-lactamase production, porin OmpK35/36 insertional inactivation, and down-regulation of the phosphate transport porin phoe, Antimicrob. Agents Chemother., 50 (2006) 3396–3406.

[11] A. Loli, L.S. Tzouvelekis, E. Tzelepi, A. Carattoli, A.C. Vatopoulos, P.T. Tassios, V. Miriagou, Sources of diversity of carbapenem resistance levels in Klebsiella pneumoniae carrying blaVIM-1, The Journal of antimicrobial chemotherapy, 58 (2006) 669–672.

[12] L. Martinez-Martinez, M.C. Conejo, A. Pascual, S. Hernandez-Alles, S. Ballesta, E. Ramirez De Arellano-Ramos, V.J. Benedi, E.J. Perea, Activities of imipenem and cephalosporins against clonally related strains of Escherichia coli hyperproducing chromosomal beta-lactamase and showing altered porin profiles, Antimicrob. Agents Chemother., 44 (2000) 2534–2536.

[13] A. Mena, V. Plasencia, L. Garcia, O. Hidalgo, J.I. Ayestaran, S. Alberti, N. Borrell, J.L. Perez, A. Oliver, Characterization of a large outbreak by CTX-M-1-producing Klebsiella pneumoniae and mechanisms leading to in vivo carbapenem resistance development, J Clin Microbiol, 44 (2006) 2831–2837.

[14] J.M. Slauch, S. Garrett, D.E. Jackson, T.J. Silhavy, EnvZ functions through OmpR to control porin gene expression in Escherichia coli K-12, J Bacteriol, 170 (1988) 439–441.

[15] L.C. Wang, L.K. Morgan, P. Godakumbura, L.J. Kenney, G.S. Anand, The inner membrane histidine kinase EnvZ senses osmolality via helix-coil transitions in the cytoplasm, EMBO J, 31 (2012) 2648–2659.

[16] Y.H. Foo, Y. Gao, H. Zhang, L.J. Kenney, Cytoplasmic sensing by the inner membrane histidine kinase EnvZ, Progress in biophysics and molecular biology, 118 (2015) 119–129.

[17] S. Chakraborty, R.S. Winardhi, L.K. Morgan, J. Yan, L.J. Kenney, Non-canonical activation of OmpR drives acid and osmotic stress responses in single bacterial cells, Nature communications, 8 (2017) 1587.

[18] H. Gerken, E.S. Charlson, E.M. Cicirelli, L.J. Kenney, R. Misra, MzrA: a novel modulator of the EnvZ/OmpR two-component regulon, Mol Microbiol, 72 (2009) 1408–1422.

[19] H. Gerken, R. Misra, MzrA-EnvZ interactions in the periplasm influence the EnvZ/OmpR two-component regulon, J Bacteriol, 192 (2010) 6271–6278.

[20] M.J. Casadaban, S.N. Cohen, Analysis of gene control signals by DNA fusion and cloning in Escherichia coli, J Mol Biol, 138 (1980) 179–207.

[21] A. Siryaporn, M. Goulian, Cross-talk suppression between the CpxA-CpxR and EnvZ-OmpR two-component systems in E. coli, Mol Microbiol, 70 (2008) 494–506.

[22] E. Batchelor, T.J. Silhavy, M. Goulian, Continuous control in bacterial regulatory circuits, J Bacteriol, 186 (2004) 7618–7625.

[23] M.H. Norholm, G. von Heijne, R.R. Draheim, Forcing the Issue: Aromatic Tuning Facilitates Stimulus-Independent Modulation of a Two-Component Signaling Circuit, ACS Synth Biol, 4 (2015) 474–481.

[24] W. Hsing, T.J. Silhavy, Function of conserved histidine-243 in phosphatase activity of EnvZ, the sensor for porin osmoregulation in Escherichia coli, J Bacteriol, 179 (1997) 3729–3735.

[25] B.J. Cantwell, R.R. Draheim, R.B. Weart, C. Nguyen, R.C. Stewart, M.D. Manson, CheZ phosphatase localizes to chemoreceptor patches via CheA-short, J Bacteriol, 185 (2003) 2354–2361.

[26] J.A. Southern, D.F. Young, F. Heaney, W.K. Baumgartner, R.E. Randall, Identification of an epitope on the P and V proteins of simian virus 5 that distinguishes between two isolates with different biological characteristics, The Journal of general virology, 72 (Pt 7) (1991) 1551–1557.

[27] M. Bogdanov, W. Zhang, J. Xie, W. Dowhan, Transmembrane protein topology mapping by the substituted cysteine accessibility method (SCAM(TM)): application to lipid-specific membrane protein topogenesis, Methods, 36 (2005) 148–171.

[28] A. Heininger, R. Yusuf, R. Lawrence, R.R. Draheim, Identification of transmembrane helix 1 (TM1) surfaces important for EnvZ dimerisation and signal output, Biochim. Biophys. Acta, 1858 (2016) 1868–1875.

[29] T. Hessa, N.M. Meindl-Beinker, A. Bernsel, H. Kim, Y. Sato, M. Lerch-Bader, I. Nilsson, S.H. White, G. von Heijne, Molecular code for transmembrane-helix recognition by the Sec61 translocon, Nature, 450 (2007) 1026–1030.

[30] A. Krogh, B. Larsson, G. von Heijne, E.L. Sonnhammer, Predicting transmembrane protein topology with a hidden Markov model: application to complete genomes, J Mol Biol, 305 (2001) 567–580.

[31] T.K. Nyholm, S. Ozdirekcan, J.A. Killian, How protein transmembrane segments sense the lipid environment, Biochemistry, 46 (2007) 1457–1465.

[32] C. Monzel, G. Unden, Transmembrane signaling in the sensor kinase DcuS of Escherichia coli: A long-range piston-type displacement of transmembrane helix 2, Proc Natl Acad Sci U S A, 112 (2015) 11042–11047.

[33] J.H. Miller, A Short Course in Bacterial Genetics: A Laboratory Manual and Handbook for Escherichia coli and Related Bacteria, Cold Spring Harbor Laboratory Press, Plainview, NY, 1992.

[34] F.M. Ausubel, R. Brent, R.E. Kingston, D.D. Moore, J.G. Seidman, J.A. Smith, K. Struhl, Current Protocols in Molecular Biology, Wiley, New York, NY, 1998.

[35] C.A. Schneider, W.S. Rasband, K.W. Eliceiri, NIH Image to ImageJ: 25 years of image analysis, Nat Methods, 9 (2012) 671–675.

[36] H. Yaku, T. Mizuno, The membrane-located osmosensory kinase, EnvZ, that contains a leucine zipper-like motif functions as a dimer in Escherichia coli, FEBS Lett, 417 (1997) 409–413.

[37] S. Forst, D. Comeau, S. Norioka, M. Inouye, Localization and membrane topology of EnvZ, a protein involved in osmoregulation of OmpF and OmpC in Escherichia coli, J Biol Chem, 262 (1987) 16433–16438.

[38] G. von Heijne, Y. Gavel, Topogenic signals in integral membrane proteins, Eur J Biochem, 174 (1988) 671–678.

[39] G. Heijne, The distribution of positively charged residues in bacterial inner membrane proteins correlates with the trans-membrane topology, EMBO J, 5 (1986) 3021–3027.

[40] I. Gushchin, I. Melnikov, V. Polovinkin, A. Ishchenko, A. Yuzhakova, P. Buslaev, G. Bourenkov, S. Grudinin, E. Round, T. Balandin, V. Borshchevskiy, D. Willbold, G. Leonard, G. Buldt, A. Popov, V. Gordeliy, Mechanism of transmembrane signaling by sensor histidine kinases, Science, (2017).

[41] E. Hwang, H.K. Cheong, S.Y. Kim, O. Kwon, K.Y. Blain, S. Choe, K.J. Yeo, Y.W. Jung, Y.H. Jeon, C. Cheong, Crystal structure of the EnvZ periplasmic domain with CHAPS, FEBS Lett, 591 (2017) 1419–1428.

[42] M. Motz, K. Jung, The role of polyproline motifs in the histidine kinase EnvZ, PloS one, 13 (2018) e0199782.

[43] K.G. Hicks, S.P. Delbecq, E. Sancho-Vaello, M.P. Blanc, K.K. Dove, L.R. Prost, M.E. Daley, K. Zeth, R.E. Klevit, S.I. Miller, Acidic pH and divalent cation sensing by PhoQ are dispensable for systemic salmonellae virulence, eLife, 4 (2015) e06792.

[44] S.S. Yadavalli, T. Goh, J.N. Carey, G. Malengo, S. Vellappan, B.E. Nickels, V. Sourjik, M. Goulian, J. Yuan, Functional determinants of a small protein controlling a broadly conserved bacterial sensor kinase, J Bacteriol, (2020).

[45] R. Yusuf, T.L. Nguyễn, A. Heininger, R.J. Lawrence, B.A. Hall, R.R. Draheim, *In vivo* cross-linking and transmembrane helix dynamics support a bidirectional non-piston model of signaling within *E. coli* EnvZ, bioRxiv, (2018) 206888.

[46] S. Kitanovic, P. Ames, J.S. Parkinson, Mutational analysis of the control cable that mediates transmembrane signaling in the Escherichia coli serine chemoreceptor, J Bacteriol, 193 (2011) 5062–5072.

[47] S. Kitanovic, P. Ames, J.S. Parkinson, A Trigger Residue for Transmembrane Signaling in the Escherichia coli Serine Chemoreceptor, J Bacteriol, 197 (2015) 2568–2579.

[48] G.A. Wright, R.L. Crowder, R.R. Draheim, M.D. Manson, Mutational analysis of the transmembrane helix 2-HAMP domain connection in the Escherichia coli aspartate chemoreceptor tar, J Bacteriol, 193 (2011) 82–90.

[49] S. Unnerstale, L. Maler, R.R. Draheim, Structural characterization of AS1-membrane interactions from a subset of HAMP domains, Biochim Biophys Acta, 1808 (2011) 2403–2412.

[50] C.A. Adase, R.R. Draheim, M.D. Manson, The residue composition of the aromatic anchor of the second transmembrane helix determines the signaling properties of the aspartate/maltose chemoreceptor Tar of Escherichia coli, Biochemistry, 51 (2012) 1925–1932.

[51] C.A. Adase, R.R. Draheim, G. Rueda, R. Desai, M.D. Manson, Residues at the cytoplasmic end of transmembrane helix 2 determine the signal output of the TarEc chemoreceptor, Biochemistry, 52 (2013) 2729–2738.

